# Seipin traps triacylglycerols to facilitate their nanoscale clustering in the ER membrane

**DOI:** 10.1101/2020.10.26.355065

**Authors:** Xavier Prasanna, Veijo T. Salo, Shiqian Li, Katharina Ven, Helena Vihinen, Eija Jokitalo, Ilpo Vattulainen, Elina Ikonen

## Abstract

Seipin is a disk-like oligomeric ER protein important for lipid droplet (LD) biogenesis and triacylglycerol (TAG) delivery to growing LDs. Here we show through biomolecular simulations bridged to experiments that seipin can trap TAGs in the ER bilayer via the luminal hydrophobic helices of the protomers delineating the inner opening of the seipin disk. This promotes the nanoscale sequestration of TAGs at a concentration that by itself is insufficient to induce TAG clustering in a lipid membrane. We identify Ser166 in the α3 helix as a favored TAG occupancy site and show that mutating it compromises the ability of seipin complexes to sequester TAG *in silico* and to promote TAG transfer to LDs in cells. While seipin-S166D mutant colocalizes poorly with promethin, the association of nascent wild-type seipin complexes with promethin is promoted by TAGs. Together, these results suggest that seipin traps TAGs via its luminal hydrophobic helices, serving as a catalyst for seeding the TAG cluster from dissolved monomers inside the seipin ring, thereby generating a favorable promethin binding interface.

## Introduction

Lipid droplets (LDs) are intracellular storage organelles critical for energy metabolism [1,2]. LDs consist of neutral lipids, mainly triglycerides (TAGs) and sterol esters, surrounded by a phospholipid (PL) monolayer. LD biogenesis takes place in the endoplasmic reticulum (ER), where DGAT enzyme activity induces TAG deposition in the bilayer [3–5]. However, PL bilayers can only accommodate low (circa 3 mol%) of TAGs before TAGs tend to phase separate. Upon rising local TAG concentration in the membrane, TAGs are thought to cluster and form nm-sized lens-like structures [6] that will then bud out to the cytoplasmic side of the ER [7,8]. The formed LDs retain a functional connectivity with the ER via membranous bridges that allow their further growth [9,10].

*In silico* work complementing experiments has turned out to be instrumental in deciphering nanoscale cellular processes [11,12]. In simulations, TAGs form lens-like lipid clusters inside a lipid bilayer, giving rise to blister-like membrane regions [13]. Their emergence from the ER can be modulated by ER topology, membrane asymmetry, surface tension and PL composition [14–19]. Moreover, the process by which the membrane properties of the TAG-engulfed LD monolayer regulate protein recruitment to LDs is starting to emerge [20–22]. However, it is not known if the initial process of TAG clustering in the ER is modulated by proteins.

The ER protein seipin is critical for LD assembly and adipogenesis [23–26], and its mutations result in lipodystrophies or neuronal disorders [27–30]. In the ER, seipin forms disk-like oligomers consisting of 11 subunits in human cells [31]. Based on cryo-EM structures of the seipin luminal regions, each protomer contains a lipid binding domain and a hydrophobic, membrane-anchored helix [31,32]. Indeed, seipin has been proposed to control cellular PL metabolism, especially phosphatidic acid handling [33] and/or to play a structural role at LD forming subdomains and ER-LD contacts [34–38].

Seipin can determine where LDs form [35]. Without seipin, LDs appear to spontaneously “oil-out” of the ER in the form of tiny and supersized LDs [34,39,40]. Seipin interacts with promethin, a multi-transmembrane protein with homology to ER-shaping reticulon proteins [41–43], and this interaction is somehow important during early LD assembly [44]. Upon LD growth promethin disassembles from seipin and decorates the surface of LDs, whilst seipin remains at the ER-LD neck. We recently reported that seipin facilitates TAG delivery to the LD and counteracts ripening-induced shrinkage of small LDs [35]. How this seipin-mediated TAG flux may be achieved, is not known. Here, we address this question by combining biomolecular simulations and cell-based experiments. Our results suggest that the seipin complex, via its hydrophobic membrane intercalated helices, can trap TAGs in the bilayer, thereby serving as a seed for TAG clustering inside the lumen of the seipin ring. This appears to generate a favorable interface for promethin binding and constitute a critical step for further LD growth.

## Results and Discussion

### TAG clusters in the lumen of seipin oligomer

To investigate the role of seipin in regulating TAG distribution, we conducted molecular dynamics (MD) simulations. We constructed a model of the seipin oligomer based on the cryo-EM structure of the luminal domains of human seipin [31] and modeled the transmembrane domains (TMDs) on the structure based on their amino acid sequence. The resulting model was placed in a multi-component bilayer, whose lipid composition mimics that of the ER **(Fig S1A, Table 1)**.

**Table 1:** Lipid composition of the membrane used in seipin oligomer simulations. The membrane was comprised of 13 different lipid species with molar concentrations identified in the Table.

| Lipid type | Total molar concentration |  |
| --- | --- | --- |
|  | *1.25 mol% TAG* | *2.5 mol% TAG* |
| 16:0-18:1 (Δ9-cis) PC | 24.27 | 23.02 |
| 18:1 (Δ9-cis) PC | 10.89 | 10.89 |
| 16:0-18:2 (Δ9-cis, Δ12-cis) PC | 7.34 | 7.34 |
| 16:0-16:1 (Δ9-cis) PC | 6.44 | 6.44 |
| 16:0-18:1 (Δ9-cis) PE | 15.03 | 15.03 |
| 18:1 (Δ9-cis) PC | 15.03 | 15.03 |
| 16:0-18:1 (Δ9-cis) PI | 10.00 | 10.00 |
| 16:0-18:1 (Δ9-cis) PS | 3.95 | 3.95 |
| 18:1 (Δ9-cis) PC | 2.95 | 2.95 |
| 16:0-18:1 (Δ9-cis) PA | 0.95 | 0.95 |
| 18:1 (Δ9-cis) PA | 0.95 | 0.95 |
| Cholesterol | 0.95 | 0.95 |
| TAG (triolein) | 1.25 | 2.50 |

An initial trigger for LD biogenesis is the generation of TAGs within the ER. We thus placed free TAG molecules (2.5 mol%) within the model bilayer with or without the seipin oligomer, and carried out coarse-grained simulations for up to 30 µs. In both systems, TAGs rapidly clustered to form lens-like structures (**Fig 1A-B)**, in line with previous *in vitro* and *in silico* work, indicating limited solubility of TAG in PL bilayers [13,45]. Remarkably, in the presence of seipin, TAG clustering occurred invariably within the lumen of the disk-shaped seipin oligomer (**Fig 1B)**.

**Fig 1.**
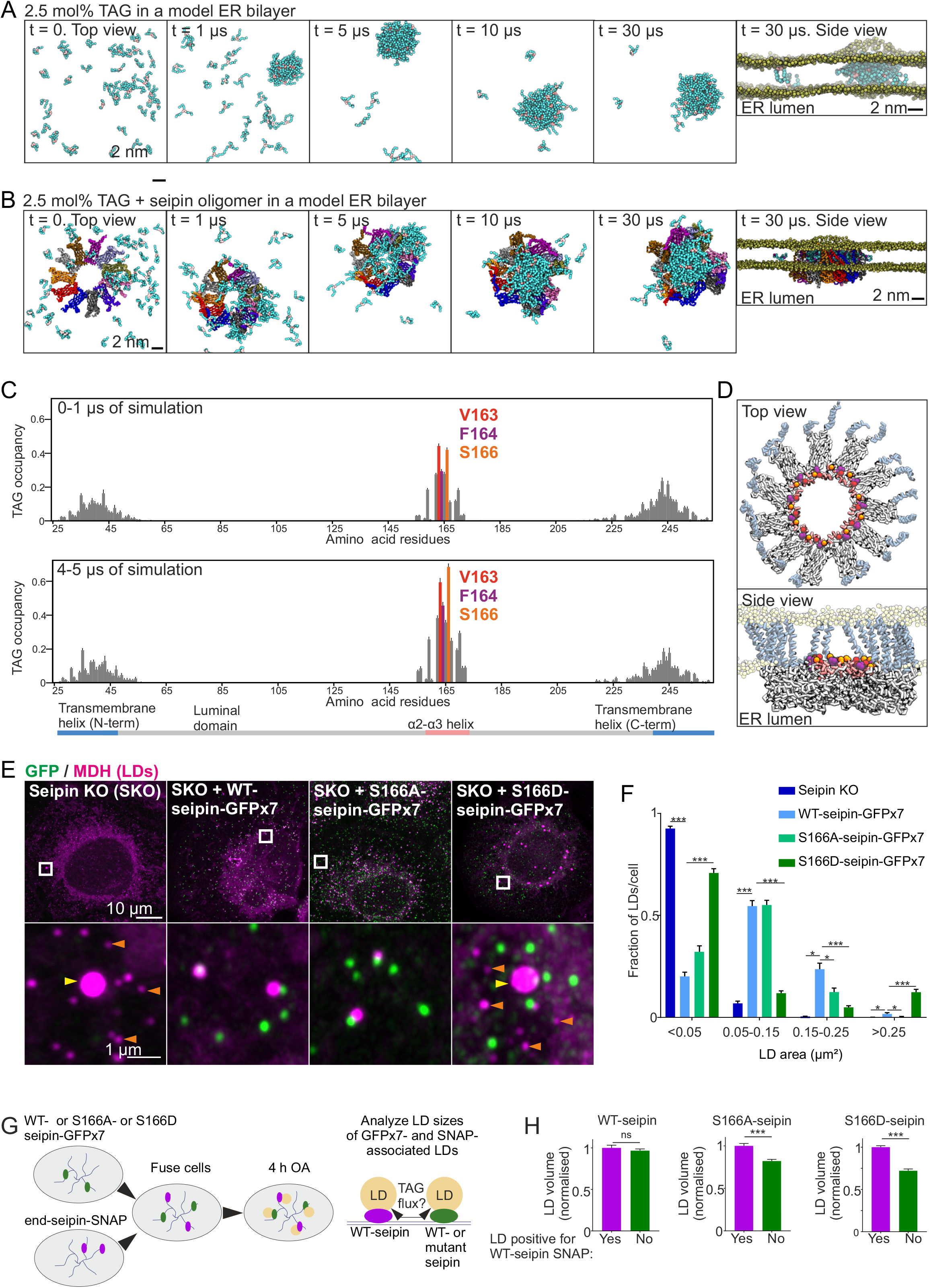
The luminal hydrophobic helix is important for TAG clustering within the seipin disk. **A)** Snapshots of coarse-grained simulations in the absence of seipin with 2.5 mol% TAG in an ER bilayer. For clarity, PL headgroups (yellow) are only shown in the side view and water is not shown. The acyl chains of TAGs are shown in cyan, glycerol moiety is in light brown. **B)** Snapshots of simulations as in A), but in the presence of seipin. **C)** TAG occupancy data of simulations demonstrated in B). TAG occupancy for each amino acid residue is defined as the probability that the residue has a TAG molecule within 0.5 nm distance from its surface. The TAG occupancies for the initial (0-1 µs) and later (4-5 µs) stages of the simulation are plotted separately. Bars = mean +/- SEM, n = 10 replicates. **D)** Key residues from C) are highlighted. **E)** A431 SKO cells stably expressing indicated plasmids were delipidated for 3 days and treated with 200 µM OA for 1 h. Cells were fixed, LDs stained, and imaged by Airyscan microscopy. Maximum intensity projections of z-stacks. Orange arrowheads: tiny LDs, yellow arrowheads: supersized LDs. **F)** Analysis of E). Bars = mean +/- SEM, n = ≥60 cells/group, 3-4 experiments. Statistics: Kruskal-Wallis test followed by Dunn’s test, comparing against WT-seipin-GFPx7. **G)** End-seipin-SNAP cells and SKO cells stably expressing WT-, S166A- or S166D-seipin-GFPx7 were co-plated for 2 days in delipidation conditions. Cells were fused with PEG and 12-14 h later 200 µM OA and Cell-SIR647 were added to the cells. 4 h after this, LDs were stained with MDH and fused cells were imaged live. **H)** Analysis of G. The sizes of end-seipin-SNAP associated LDs were compared to LDs within the same cell not positive for SNAP. Bars = mean +/- SEM. n = 152-659 LDs/group, 4-20 fused cells/group, 2 experiments. Statistics: Mann-Whitney test. Exemplary micrographs are shown in Fig S1D.

We next mapped the TAG occupancy of seipin amino acid residues by analyzing the probability of each residue to be in close proximity to a TAG molecule. The TMDs showed affinity for TAG but the highest TAG occupancy, especially at later simulation times, was observed for three residues in the hydrophobic, membrane-anchored α2-α3 helix of seipin, namely V163, F164 and S166 (**Fig 1C-D**). These data suggest that TAGs cluster in the lumen of the seipin oligomer and that distinct residues of the hydrophobic helices delineating the inner opening of the seipin ring play an important role in this.

### S166 is critical for seipin function

To assess the relevance of the residues identified as preferential TAG interaction sites, we analyzed whether mutating them would affect seipin function in LD biogenesis. We designed a panel of single amino acid substitution mutants, tagged them with split 7xGFP [46], and expressed in human A431 seipin knockout (SKO) cells. We used FACS to enrich stable pools with near-endogenous seipin expression levels. Cells were then delipidated, treated with oleic acid (OA) to induce LD biogenesis, and imaged by Airyscan microscopy. In cells without detectable GFP (*i.e*. SKO cells), LDs were tiny or supersized, as expected **(Fig 1E-F)**. Expression of WT-seipin-GFPx7 rescued this phenotype, with single seipin foci associated with virtually all LDs (**Fig 1E-F, S1B**), as previously reported [34]. Mutations affecting V163 or F164 led to a more diffuse reticular GFP fluorescence pattern compared to WT-seipin, suggesting they may affect seipin complex assembly **(Fig S1B)**. However, single seipin foci could also still be discerned and these were often juxtaposed to LDs, and the LD morphology was similar to that of the WT-seipin-GFP expressing cells (**Fig S1B)**. The notion that the oligomerization propensity of seipin may be sensitive to mutations in V163 and F164 is in line with the fact that they reside at the seipin protomer-protomer interface [31].

In contrast to mutations affecting V163 and F164, mutation of S166 did not appear to impair seipin complex formation, as the mutants tested existed as discrete puncta in cells (**Fig S1B**). S166A-seipin rescued the overall LD morphology in SKO, but LDs were on average somewhat smaller than in WT-seipin expressing cells **(Fig 1E-F, S1B-C)**. Importantly, S166D-seipin was not able to complement the LD biogenesis defect in SKO, as most cells contained supersized and tiny LDs **(Fig 1E-F, S1B-C)**.

We previously reported that seipin can promote the growth of its associated LD: in a continuous ER with both seipin-containing LDs and LDs where seipin had been acutely depleted, seipin containing LDs grew at the expense of seipin depleted ones [35]. To study whether the S166-mutants can promote the growth of existing LDs as well as WT-seipin, we performed competition experiments employing heterologous cell fusions (**Fig 1G)**. In a continuous ER network with LDs associated with either S166-mutant or WT-seipin, the growth of S166D- or S166A-seipin associated LDs was reduced compared to WT-seipin associated LDs, suggesting that the S166-mutants are defective in promoting LD growth (**Fig 1G-H, S1D)**. Altogether, these data imply that mutating S166 to Asp impairs seipin function both in initial LD assembly and subsequent LD growth, whilst mutating S166 to Ala affects the growth of assembled LDs.

Having identified a key seipin residue for TAG interaction, we performed atomistic MD simulations of the situation where TAGs are clustered within the seipin disk. These revealed that S166 tends to form a complex with TAG (**Fig 2A-B**). Simultaneous interaction was observed between a TAG molecule and S166 as well as the neighboring S165 (**Fig 2 B**). The interaction involved formation of a hydrogen bond between a carbonyl group of the TAG molecule with the amino acid backbone of S166. This interaction appeared to be further stabilized by formation of a transient hydrogen bond between another carbonyl group of the same TAG molecule with the side chain of S165, reducing the diffusion of the associated TAG. By calculating the short-time (ns scale) diffusion coefficient *D*, we observed that the *D* of TAGs not interacting with S166 (magenta trajectory in **Fig 2A**) was ∼3 times larger than the *D* of TAG bound to S166 (red trajectory in **Fig 2A**).

**Fig 2.**
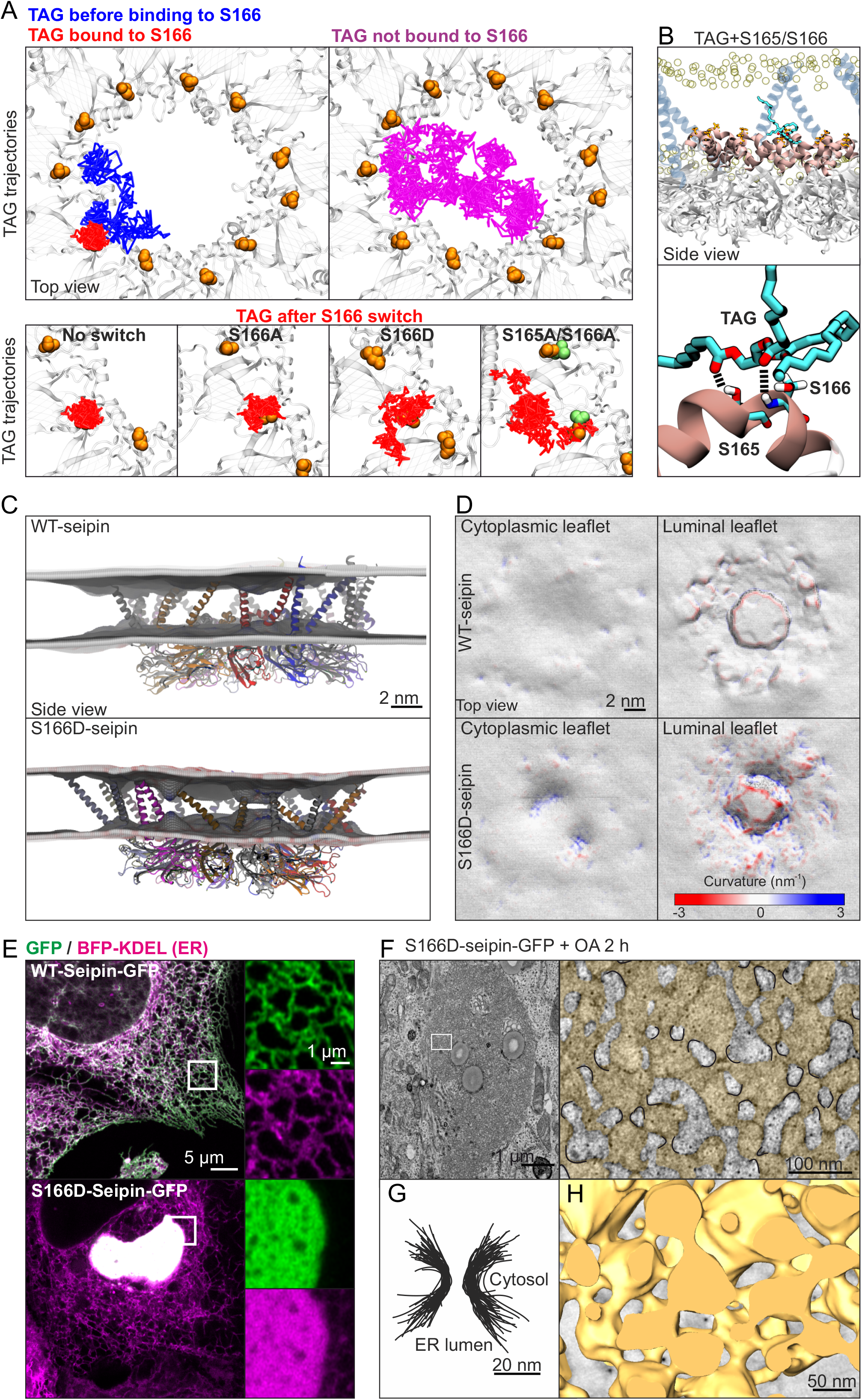
S166D-seipin impairs TAG interaction and induces ER membrane deformation. **A)** Exemplary TAG trajectories from an atomistic simulation with a TAG cluster within the seipin disk. Blue indicates TAG trajectory prior to S166 binding (0-590 ns), red shows its trajectory upon S166 binding (591-960 ns). Magenta indicates the trajectory of TAG not interacting with S166 (0-960 ns). Lower row, left: higher magnification of S166-binding TAG trajectory, other insets: TAG trajectories after indicated amino acid changes (trajectory lengths are 370 ns for all insets). Each line represents a series of in-plane 1 ns displacements. **B)** Snapshot of the interface between a TAG molecule and S165 and S166. Hydrogen bonds between the carbonyl groups of TAG molecule and S165 and S166 are represented by black dashed parallel lines. The α2-α3 helix is shown in pink. **C)** Snapshots of atomistic simulations of WT- or S166-seipin, performed with 2.5 mol% TAG in the membrane. The time-averaged (>1 μs) mean curvature of bilayer leaflets from the plane of PL headgroups is shown. **D)** Topologies and the mean curvature of the bilayer leaflets from C) is shown. **E)** A431 cells stably expressing BFP-KDEL were transfected with WT- or S166D-seipin-GFP for 3 days and imaged live by Airyscan microscopy. **E)** HEK293A cells were transfected with S166D-seipin-GFP for 3 days in delipidation conditions and treated with 200 µM OA for 2 h. TEM images of the membranous aggregates display periodical ER constrictions (indicated with black in the zoom-in, ER is colored with yellow). **F)** Segmented membrane constrictions from F) are overlaid. **G)** Cells were treated as in F and imaged by electron tomography. A 3D model of the ER sacs (yellow) induced by S166D-seipin-GFP is shown.

To simulate S166-mutants in this context, we changed all S166 residues to either Asp or Ala after the formation of a stable S166-TAG complex. The simulations revealed that interaction of TAG with S166D was significantly weaker than with S166, whilst switching to S166A did not similarly impair the interaction. The short-time diffusion coefficient of TAG bound to S166D was ∼50% larger than that bound to S166 (**Fig 2A**). The S166D-TAG complexes were also less stable, with a life time of ∼0.1 μs compared to ∼0.5 μs of S166-TAG and S166A-TAG complexes. The short-time diffusion coefficient of TAG bound to S166A was ∼15% larger than that bound to S166. Finally, as 165A/166A double mutant has been reported to impair seipin function [44], we switched both S165 and S166 to Ala and found that this switch also impaired TAG-S166 interaction, with the short-time diffusion coefficient of TAG being ∼100% larger than the *D* of TAG bound to S166, and a life time of ∼0.1 µs for the S166A-TAG complex.

In summary, these data suggest that S166 serves as a nucleation site in the formation of TAG clusters. Given that there are eleven S166 residues in the seipin complex, it seems evident that the TAG clusters forming around S166 residues gradually merge and promote the growth of a larger cluster.

### S166D-seipin overexpression induces ER membrane deformation

We next assessed the effects of the S166D change on the neighboring PLs in simulations with or without a TAG cluster in the seipin disk. There was no significant movement of the α2-α3 helices towards the core of the bilayer in either condition in WT- or S166D-seipin containing systems (**Fig S2A-D)**. However, the bilayer was locally deformed in the presence of S166D-seipin, with apparent bulging of the PL head groups towards the membrane interior on both bilayer leaflets (**Fig 2C**). In line with this, topological maps of the leaflets displayed higher negative and positive curvatures in simulations with S166D-compared to WT-seipin **(Fig 2D)**. These effects were apparently related to the negative charge of Asp, as protonation of S166D in the simulations alleviated the membrane perturbation (**Fig S2E-F)**.

In cells stably expressing S166D-seipin, the mutant protein was present as typical seipin puncta (**Fig 1E**). However, higher transient overexpression of S166D-seipin induced a gradual, dramatic reorganization of the ER into membranous aggregates, whilst high expression of WT-seipin did not induce this (**Fig 2E, S2G**). EM analysis revealed that these aggregates consisted of small ER sacs connected by membranous constrictions **(Fig 2F-H)**. Intriguingly, these constrictions were ∼15 nm in diameter, similarly as the seipin oligomer disk. It is tempting to speculate that the bilayer destabilization observed with Asp in simulations may contribute to these ER membrane deformations, although the protonation status of S166D in cells remains open.

### S166D-seipin shows reduced colocalization with promethin

The interaction of promethin and seipin requires the luminal hydrophobic helix of seipin [44], which also harbors the amino acids identified in our simulations as maximal TAG occupancy sites. We thus reasoned that the S166-mutants may impact seipin-promethin interaction. To investigate this, we performed immunofluorescence stainings using anti-promethin antibodies. These reliably detected endogenous promethin, as evidenced by using CRISPR/Cas9 engineered promethin KO cells as controls (**Fig S3A-B**). Furthermore, in line with previous reports [41,44], the antibodies demonstrated that promethin levels increased upon prolonged OA incubation and this was accompanied by relocalization of promethin to rings around LDs (**Fig S3C-E**). In cells grown in complete medium, we found promethin as discrete dots, many of which overlapped with WT-seipin dots (**Fig 3A-B)**. Indeed, ∼60% of WT-seipin foci were positive for promethin and similar results were found for S166A-seipin **(Fig 3A-B, S3F)**. In contrast, only a small fraction (∼10%) of S166D-seipin overlapped with promethin, indicating that S166D impairs seipin association with promethin (**Fig 3A-B, S3F)**.

**Fig 3.**
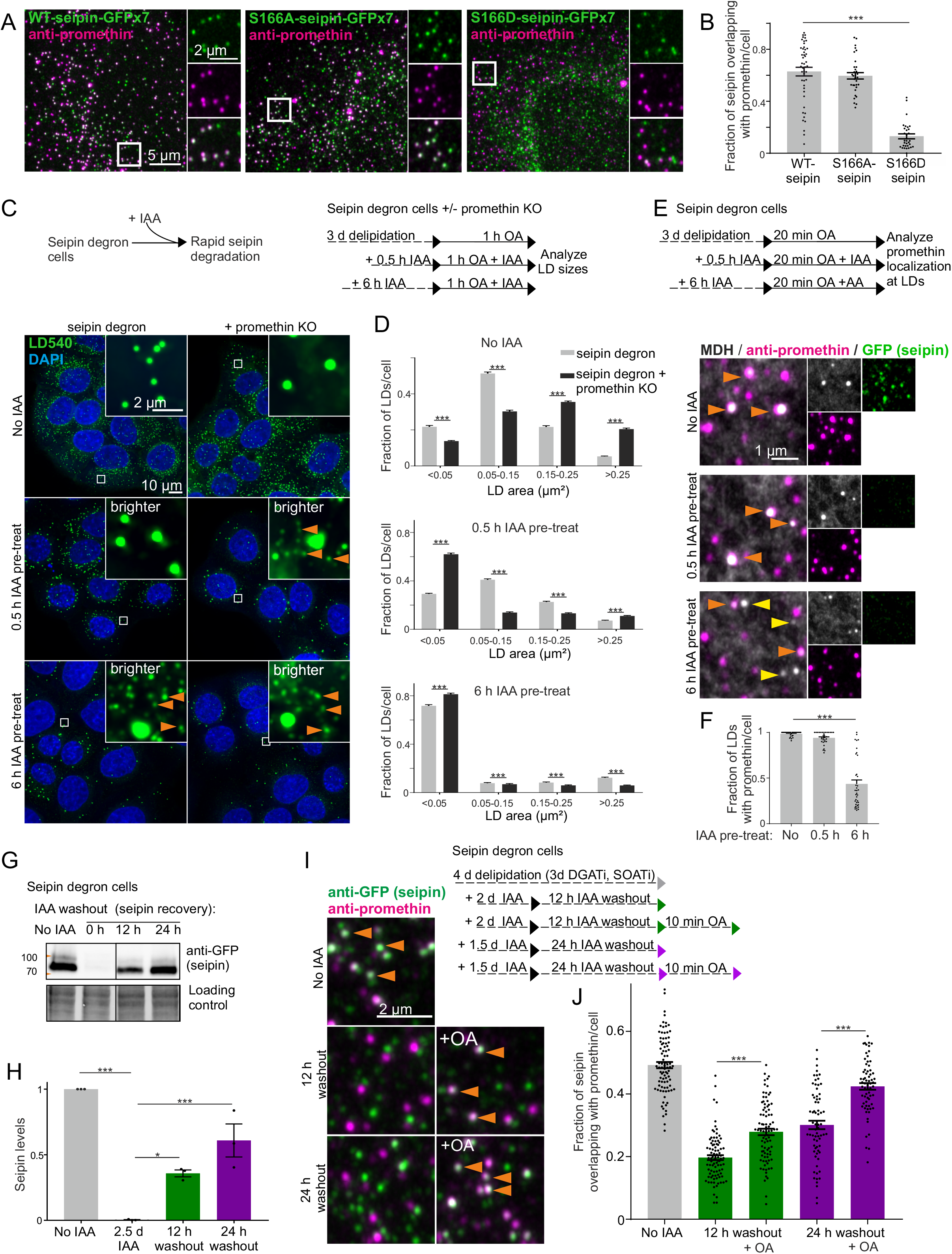
Promethin sensitizes cells to seipin depletion and TAG facilitates nascent seipin association with promethin. **A)** A431 SKO cells stably expressing indicated plasmids were fixed and stained with anti-promethin antibodies. **B)** Analysis of A. Bars = mean +/- SEM, n = ≥30 cells/group, 3-4 experiments. Statistics: Kruskal-Wallis test followed by Dunn’s test, comparing to WT-seipin. **C)** Seipin degron cells with or without promethin KO were delipidated for 3 d, treated with IAA and 200 µM OA as indicated, fixed, stained, and imaged by widefield microscopy. Deconvolved maximum intensity projections of z-stacks. Orange arrowheads: tiny LDs. Note that 0.5 h IAA pre-treatment is sufficient to induce generation of tiny LDs in promethin KO cells. **D)** Analysis of C). Bars = mean +/- SEM, n = >500 cells/group, 3 experiments. Promethin KO data are pooled from two KO pools. Statistics: Mann-Whitney test. **E)** Seipin degron cells were delipidated for 3 d, including the last 18 h in the presence of DGAT inhibitors. Cells were treated with IAA as indicated. The inhibitors were washed out and cells were treated with 200 µM OA, fixed, stained with anti-promethin antibodies and MDH, and imaged by Airyscan microscopy. Crops of maximum intensity projections of Airyscan z-stacks. Orange arrowheads: LDs with promethin signal; yellow arrowheads: LDs without promethin signal. **F)** Analysis of E. Bars = mean +/- SEM, n = ≥29 cells/group, 2 experiments. Statistics: Kruskal-Wallis test followed by Dunn’s test, comparing to no IAA treatment. **G)** Immunoblot of seipin levels in seipin degron cells after IAA washout as indicated. Note the recovery of seipin levels upon IAA washout. **H)** Analysis of G). Bars = mean +/- SEM, n = 3 biological replicates from 2 experiments. Statistics: 1-way ANOVA followed by Tukey’s post hoc test. **I)** Seipin degron cells were delipidated and treated with IAA and OA as indicated. DGAT and SOAT inhibitors were washed out prior to OA addition. Cells were fixed, stained with antibodies and imaged. Crops from maximum projections of Airyscan z-stacks. Orange arrowheads: seipin foci overlapping with promethin. **J)** Analysis of I. Bars = mean +/- SEM, n = >60 cells/group. Data are pooled from 4 experiments, 2 using anti-GFP and 2 using anti-degron tag antibodies for seipin detection. Statistics: Kruskal-Wallis test followed by Dunn’s test.

### Promethin depletion sensitizes cells to rapid seipin removal

To further investigate the interplay between seipin and promethin, we generated promethin KO cell pools on top of seipin degron cells, where seipin can be rapidly depleted using the auxin-inducible degron (AID) system [35,47,48]. Promethin depletion did not interfere with the AID system, as seipin was efficiently depleted in both control and promethin KO cells upon addition of indole-3-acetic acid (IAA) **(Fig S3G-H)**. Upon promethin KO, seipin levels increased **(Fig S3G-H)** and LDs were fewer in number but larger in size **(Fig S3I)**, as reported [44].

This system allowed us to scrutinize the effects of rapid seipin removal on LD biogenesis in the presence vs. absence of promethin. We have previously noted that upon acute seipin degradation (from promethin containing cells), the characteristic phenotype of numerous tiny and few supersized LDs only appears after a refractory period [35]. Thus, removing seipin only 30 min prior to LD induction is not sufficient to evoke the SKO phenotype, while it becomes apparent when LDs are induced after a longer (6 h) seipin depletion (**Fig 3C-D**). Interestingly, we found that loss of promethin sensitized cells to seipin depletion, as in promethin KO cells tiny and supersized LDs appeared when seipin was removed just 30 min prior to LD induction (**Fig 3C-D**). We also investigated the localization of promethin at nascent LDs under these conditions. In cells containing seipin, virtually all nascent LDs were positive for promethin and this was also the case if seipin had been depleted for only 30 min prior to LD induction **(Fig 3E-F)**. However, upon longer 6 h seipin depletion, a large fraction (over 50% on average) of nascent LDs generated did not contain promethin foci **(Fig 3E-F)**.

The above findings are compatible with the idea that the seipin-promethin complex may harbor miniature TAG clusters [44], even in delipidated conditions. Upon seipin removal from promethin containing cells, these structures may initially be stable and serve as assembly sites for LDs, contributing to the latency in developing the SKO LD phenotype. Upon seipin removal from promethin deficient cells, they may be destabilized more rapidly and LD biogenesis proceeds stochastically, with the hallmark of numerous tiny LDs. These data do not rule out other factors, e.g. a more general perturbation of lipid metabolism, that may contribute to the delayed development of the SKO phenotype in the presence of promethin.

### TAG facilitates seipin-promethin association

Given that S166D-seipin was dysfunctional in LD biogenesis and failed to colocalize with promethin, we reasoned that the association of TAGs with the seipin oligomer might facilitate seipin-promethin association. However, even prolonged lipid starvation in the presence of neutral lipid synthesizing enzyme inhibitors did not disassemble the promethin-seipin foci (**Fig 3 I-J)**. One possibility is that after initial assembly, promethin-seipin complexes are stable. We therefore investigated the interaction of newly formed seipin oligomers and promethin, taking advantage of seipin degron cells.

Upon IAA washout, seipin levels recovered with an apparent half-time of ∼16 h, as the cells generated new seipin protein not subjected to degradation **(Fig 3G-H)**. We therefore removed seipin in delipidating conditions (in the presence of neutral lipid synthesizing enzyme inhibitors) and allowed seipin levels to recover for 12 or 24 h (**Fig 3I-J)**. Then, we washed out the inhibitors and pulsed the cells with OA for 10 min to induce TAG synthesis. When seipin foci were synthesized under lipid poor conditions, a smaller fraction of seipin overlapped with promethin foci compared to cells where seipin had not been depleted. Remarkably, the 10 min OA pulse increased the fraction of seipin foci overlapping with promethin **(Fig 3I-J)**. Similar results were obtained when considering the fraction of promethin overlapping with seipin **(Fig S3J-K, S3K)**. Together, these data suggest that neutral lipids can facilitate the colocalization of seipin and promethin. We speculate that this may arise from local membrane alterations induced by TAG clustering within the seipin oligomer, providing a favorable environment for promethin association.

### Seipin hydrophobic helix provides a seed for TAG nucleation

Our data suggest a model in which seipin attracts TAG from the ER bilayer to its lumen, which then leads to promethin recruitment and, with rising TAG concentration, to LD formation. As our simulations using 2.5 mol% TAG resulted in TAG clustering already in the absence of seipin (**Fig 1A**), we investigated TAG clustering at a lower concentration (1.25 mol% TAG), comparing systems without seipin, with WT-seipin and with various seipin mutants. In the absence of seipin, TAG molecules now failed to cluster, whilst WT-seipin facilitated rapid redistribution of TAG to the oligomer lumen **(Fig 4A-B)**. Other transmembrane proteins tested did not efficiently catalyze TAG clustering under these conditions **(Fig S4A-C)**. S166A-seipin facilitated initial TAG clustering, but the process was compromised in the case of S166D- and S165A/S166A-seipin (**Fig 4A-B**). Interestingly, TAG clustering was also achieved by seipin oligomer without its TMDs, suggesting that TMDs are not required for this activity (**Fig S4A-B)**. These data suggest that seipin can efficiently concentrate TAG molecules from the surrounding ER bilayer to its lumen and this is hampered by S166D- or S165A/S166A-seipin.

**Fig 4.**
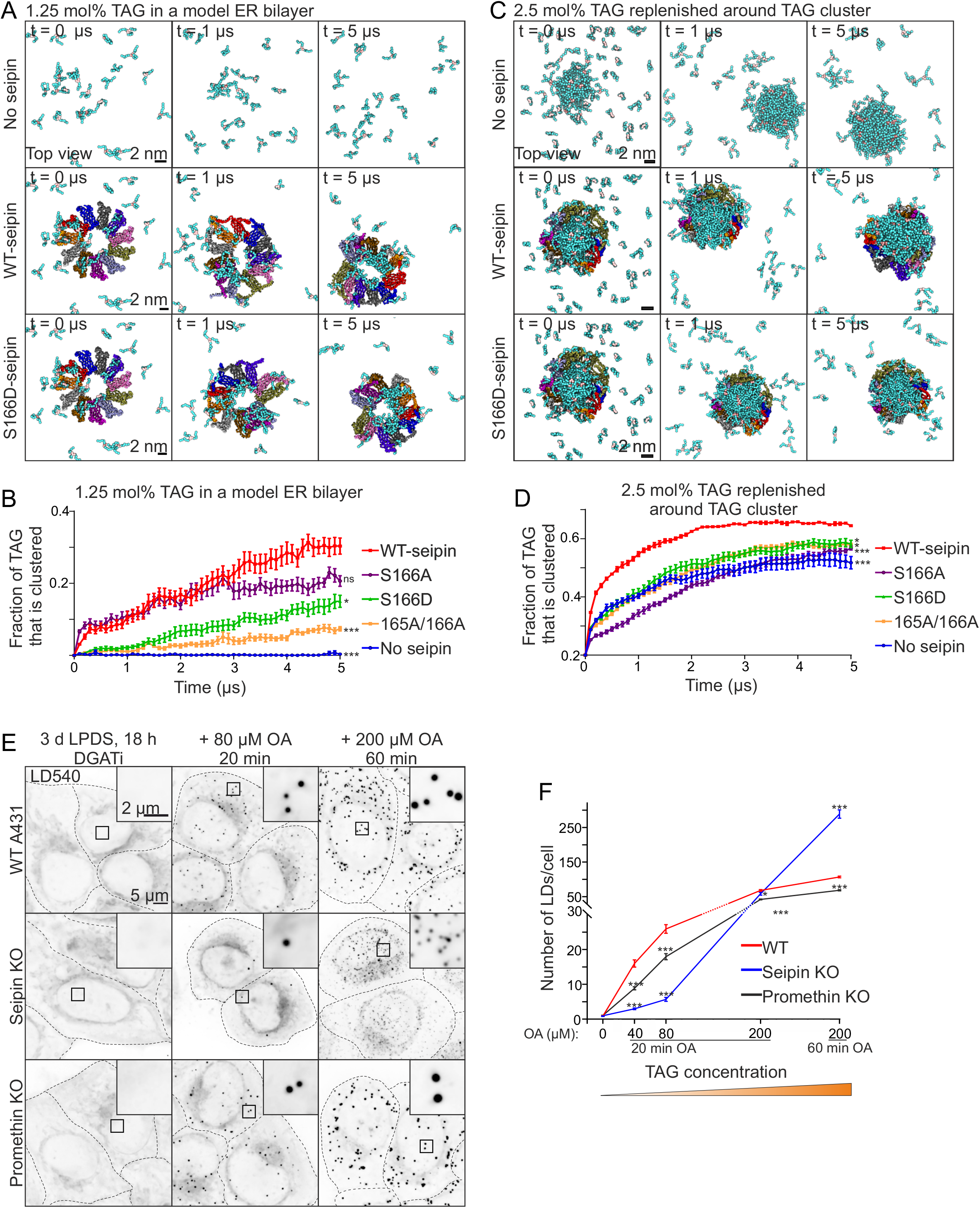
Seipin promotes the nanoscale clustering of TAGs and LD biogenesis at low TAG concentration. **A)** Snapshots of coarse-grained simulations with 1.25 mol% TAG in the bilayer. Same color coding as in Fig 1A. **B)** Analysis of simulations carried out as in A). Data points: mean, +/- SEM, n = 10 simulations/system. Statistics are based on the final time points of analysis using Kruskal-Wallis test followed by Dunn’s test, comparing to WT-seipin. **C)** Snapshots of coarse-grained simulations with an initial TAG cluster and the rest of the model bilayer replenished with 2.5 mol% TAG (total TAG concentration 4.85 mol%). **D)** Analysis of simulations carried out as in C). Data points: mean +/- SEM, n = 10 simulations/system. Kruskal-Wallis test followed by Dunn’s test, statistics based on the final time points of analysis, comparing to WT-seipin. **E)** Cells were delipidated for 3 d, including 18 h in the presence of DGAT inhibitors. The inhibitors were washed out and cells were treated with OA as indicated, fixed, stained and imaged by widefield microscopy. Deconvolved maximum intensity projections of z-stacks. **F)** Analysis of E). Data points = mean +/- SEM, n = >250 cells/group, 2 experiments. Statistics: 1-way ANOVA followed by Tukey’s post hoc test, comparison to WT cells. Note that the y-axis is discontinuous.

During LD biogenesis, new TAGs are constantly made via the activity of the ER-resident DGAT enzymes. To mimic this lipogenic condition in simulations, we used the conformation of seipin that hosted a clustered TAG lens in its lumen (30 µs end point in **Fig 1A**) as a starting point and replenished the rest of the model ER bilayer with 2.5 mol% TAG. The initially free TAG molecules rapidly started to coalesce with the pre-existing TAG cluster, and this was more rapid and efficient in the presence of WT-seipin (**Fig 4C-D)**. S166D and S165A/S166A-seipin failed to catalyze this TAG flux as efficiently as WT-seipin. Interestingly, S166A-seipin was also defective in this (**Fig 4D)**, in line with the observation that S166A was compromised in promoting LD growth (**Fig 1E-F, S1D-F)**. Together, these data imply that seipin can facilitate TAG flux from nearby ER to a pre-existing TAG cluster. Considering the localization of seipin at the ER-LD neck [35], this situation might be reminiscent of an ER-connected nascent LD that grows by additional TAG entering via the seipin disk, as proposed [35,44].

### Seipin promotes LD biogenesis at low TAG concentration

The MD simulations suggested that seipin may facilitate nanoscale clustering of TAGs at a low TAG concentration. To scrutinize this notion in cells, we treated stringently starved WT and SKO cells with increasing concentrations of OA for short periods of time and analyzed the number of forming LDs. We found that at low OA concentrations, SKO cells formed significantly less LDs than WT cells **(Fig 4E-F)**. This indicates that seipin is critical to catalyze LD formation at low TAG concentrations.

For comparison, we also performed similar experiments in promethin KO cells. At low OA concentrations, these cells also formed less LDs than WT cells **(Fig 4E-F)**, but more than SKO cells. Importantly, upon further incubation with high OA concentration, numerous tiny LDs became apparent in SKO but not in promethin KO cells **(Fig 4E-F)**. Together, these results suggest that while both seipin and promethin are important for LD formation at low TAG concentration, seipin is required to keep the ER TAG concentration sufficiently low to prevent spontaneous “oiling-out” of LDs.

To summarize, our data provide a plausible model of how seipin can attract TAG molecules in the ER. This activity depends on the fidelity of the α3 membrane-anchored helix and a critical S166 therein, with S166 mutations diminishing seipin activity both *in silico* and in cells. These residues serve as catalysts for concentrating TAGs inside the seipin ring, promoting nanoscale TAG clustering even at minute concentrations. The cylindrical multiunit architecture of seipin is important, providing multiple nucleation sites for TAG clusters. This TAG attracting propensity of seipin may explain why LDs preferentially form at seipin-defined sites in cells [35,44]. We propose that the TAG-affiliated seipin then recruits promethin, to stabilize the nascent lens through mechanisms yet to be defined. Additional TAGs continuously synthesized in the ER are attracted by seipin and tend to coalesce with the pre-existing seipin-trapped TAG nucleus. Such TAG harvesting molecular function of seipin likely contributes to its role in preventing ripening and sustaining LD growth [35].

## Materials and methods

### Materials and reagents

Concentrations in brackets indicate final concentrations used. Rabbit anti-GFP (Abcam, ab290, WB: 1:10 000), mouse anti-GFP (Abcam, ab1218, IF: 1:250), anti-degron tag (anti-miniIAA7; Ximbio, 158027, 1:500 IF), rabbit anti-TMEM159/promethin (Sigma, HPA063509, IF: 1:150, WB: 1:500); goat anti-mouse Alexa 488 (Thermo Scientific, A-11001, IF 1:400); goat anti-rabbit Alexa 560 (Thermo Scientific, A11011, IF 1:400); Monodansylpentane /MDH/Autodot (Abgent, SM1000a, 0.1 mM); LD540 [49] (synthesized by Princeton BioMolecular Research, 0.1 µg/ml), DAPI (Sigma, D9542, 10μg/ml); SNAP Cell-SIR647 (New England BioLabs, S9102S, 0.3 µM); PEG 1500 (Roche, 50% w/v in 75 mM Hepes); DGAT1 inhibitor (Sigma PZ0207, 5 µM); DGAT2 inhibitor (Sigma, PZ0233, 5 µM); SOAT inhibitor (Sigma, S9318, 2 µg/ml); Indole-3-acetic acid sodium (IAA, Santa Cruz, sc-215171, 500 µM); Human fibronectin (Roche Diagnostics, 11051407001); Puromycin (Sigma, P8833, 1 µg/ml); Geneticin (G418, Gibco, 11811-031, 0.6 mg/ml); Zeocin (Thermo Fisher, R25001, 0.2 mg/ml); formaldehyde (Sigma, P-6148 for light microscopy; Electron Microscopy Sciences, EMS-15710 for EM); ECL Clarity (Bio-Rad, 170-5060); ECL Clarity Max substrate (Bio-Rad, 170-5062); Saponin (Sigma-Aldrich S-7900); 12% Mini-Protean TGX Stain-Free gels (Bio-Rad, 161-0185); Nitrocellulose membranes (Bio-Rad, 170-4270); BioRad DC Protein assay (Bio-Rad, 5000112); 8-well Lab-Tek II #1.5 coverglass slides (Thermo Fisher, 155409); #1.5 polymer µ-slide 8-well and 16-well ibiTreat chambers (Ibidi, 80826 and 81816); HQ SILVER Enhancement kit (Nanoprobes, 2012); Glutaraldehyde (EM-grade, Sigma, G7651); Osmium tetroxide (Electron Microscopy Sciences, RT 19130); Epon (TAAB, 812). Other reagents, including cell culture reagents were from Gibco/Thermo Fisher, Lonza and Sigma, Merck and Honeywell.

### Cell culture

A431 (ATCC CRL-1555) and HEK 293A cells (Invitrogen, R70507) were maintained in Dulbecco’s modified Eagle’s medium containing 10% fetal bovine serum (FBS), penicillin/streptomycin (100 U/ml each), L-glutamine (2 mM) at 37°C 5% CO2.

### Plasmids and stable cell lines

A431 SKO cells, generated using CRISPR/Cas9 technology, have been described [34]. The clone used in this study corresponds to S2AB-15 in [34]. End-seipin-GFPx7 cells and end-seipin-SNAPf cells have been described [35]. End-seipin-SNAPf cells stably expressing BFP-KDEL have been described [19]. Seipin degron cells have been described [47]. These cells harbor miniIAA7-GFP at the C-terminus of endogenous BSCL2/seipin locus and express *At*AFB2, allowing for efficient depletion of seipin in response to IAA but minimal basal degradation in the absence of IAA.

Promethin KO cells were generated similarly as described [34]. Briefly, WT A431 cells were co-transfected with Cas9 nickase containing zeocin selection marker and two matching pairs of the sgRNA-expressing plasmids (sgRNA targets: gtttctggccttcaccttgc <u>tgg</u> and ctgtccaagtactgacccac <u>tgg</u>). 24 h after transfection, cells were placed under zeocin selection for 48 h. After zeocin selection, the promethin KO pool was cultured in complete medium in the absence of zeocin and used for experiments. For promethin KO on top of seipin degron cells, seipin degron cells were treated as described above, but two promethin KO pools were made, using either the sgRNAs described above (Promethin KO-B) as well as another pair of sgRNAs (Promethin KO-A, targets: cttggacagccatccgtttc <u>tgg</u> and acccactggagacttcataa <u>agg</u>). Both sgRNA pairs were efficient, as assessed by immunoblotting and LD phenotype.

For generating constructs overexpressing seipin with different point mutations, overlapping PCR was used to introduce the point mutations, and to link GFP11×7 tag (from Addgene Plasmid #70224, a gift from Bo Huang [46]) via a flexible linker to the C*-*terminus of seipin fragments. These fragments were inserted into pmCherry-N1 backbone through SalI and BsrGI restriction sites, resulting in the substitution of the mCherry fragment in the vector. All point mutations were verified by Sanger sequencing. For generating stable SKO cells expressing various seipin point mutants, SKO cells were transfected with pSH-EFIRES-P-GFP(1-10)opti AAVS1 Safe Harbor integration plasmid (Addgene plasmid # 129416, [35] and pCas9-sgAAVS1-2 (Addgene plasmid # 129727 [47]), and a stable pool was selected using puromycin (1 µg/ml puromycin for 6 days, then 5 µg/ml puromycin for 2 days, then cultured further without puromycin). These cells were then transfected with the various point mutant seipin-GFP11×7 constructs and stable pools were selected using 0.6 mg/ml G418. Cells were FACS-sorted using BD Influx Flow Cytometer (BD Biosciences, USA) at HiLIFE Biomedicum Flow cytometry unit, University of Helsinki to enrich cells with GFP fluorescence levels in the range of 0.8-2 fold compared to endogenously GFPx7-tagged seipin, using end-seipin-GFPx7 cells [35] as a control. Finally, single clones were isolated using limiting dilution to obtain SKO + WT-seipin-GFPx7, SKO + S166A-seipin-GFPx7 and SKO + S166D-seipin-GFPx7 cells. For transient overexpression (**Fig 2 E-H, S2G)** seipin tagged with C-terminal EGFP [34] was inserted into pcDNA5/FRT/TO (Thermo Fisher Scientific) through HindIII/NotI sites. S166D point mutation was introduced by overlap PCR.

### Transfection, lipid and IAA treatments, SNAP-labeling, Cell fusion

Transfection of plasmids was carried out using Lipofectamine LTX with PLUS Reagent or X-tremeGENE HP DNA according to manufacturer’s instructions. Cells were delipidated by culturing in serum-free medium supplemented with 5% LPDS for indicated times. Where indicated, for more stringent delipidation cells were additionally incubated with DGAT1, DGAT2 and SOAT inhibitors. DGAT inhibitors indicates both DGAT1 and DGAT2 inhibitors. These inhibitors were washed out by 4 washes with PBS. For LD induction cells were supplemented with the indicated concentration of OA (OA in complex with BSA in 8:1 molar ratio prepared in serum-free DMEM as described in [50]). For seipin depletion, IAA was added to the medium. For IAA washout, cells were washed with PBS for 4 times and cultured in a medium without IAA. SNAP-labeling was done for 5 min at +37°C, the label Cell-SIR647 was applied in n 5% LPDS containing medium followed by 3 washes with PBS. SNAP-labeling was performed 4 h prior to imaging experiments. Cells were fused with PEG1500 for 1 min, followed by 4 washes with PBS and further incubation in complete medium with OA.

### Immunofluorescence staining

Immunofluorescence was performed as described [35], using 0.1% saponin for permeabilization and 10% FBS for blocking and antibody dilutions.

### Immunoblotting

Cells were lysed into RIPA-buffer (1% Igepal CA-630, 0.1% SDS in TBS) containing protease inhibitors. After lysis, lysate was cleared by centrifugation at 16 000 G for 10 min at +4° C and the supernatant was collected. Protein concentrations were measured using DC protein assay. For anti-promethin immunoblotting, samples in 1X Laemmli buffer were incubated at RT for 15 min, heated at +70° C for 5 min and cooled down at RT for 30 min prior to loading onto gels. For anti-GFP immunoblotting, samples in 1X Laemmli buffer were kept on ice prior to loading onto gels without heating. Equal amounts of protein were loaded onto 12% Mini-Protean TGX Stain-Free gels and transferred onto nitrocellulose membrane. Membranes were blocked with 5% milk in TBS containing 0.1% Tween-20 for 1 h at RT, and probed with primary antibodies at +4° C overnight. After washing with TBS containing 0.1% Tween-20, membranes were incubated with secondary antibodies for 1-2 h at RT. Membranes were washed, incubated with ECL or ECL Clarity Max substrate, and imaged with a ChemiDoc Imaging System (Bio-Rad). Band intensities were quantified in ImageJ FIJI and normalized to total protein content quantified with the Stain-Free technology (Bio-Rad). Stain-Free total protein signals are shown as loading controls in the figures.

### Light microscopy and image analysis

#### Live and fixed cell imaging

Cells were seeded onto Ibidi µ-slide 8 or 18-well ibiTreat chambers or #1.5 coverglasses for widefield microscopy; or 8-well Lab-Tek II #1.5 coverglass slides for Airyscan microscopy, the latter coated with 10 mg/ml fibronectin. All live cell imaging experiments were performed at +37° C, 5% CO2 in FluoroBrite DMEM supplemented with 10% FBS. For light microscopy of fixed cells, cells were washed with PBS, fixed with 4% PFA in 250 mM Hepes, pH 7.4, 100 mM CaCl2 and 100 mM MgCl2 for 20 min, followed by quenching in 50 mM NH4Cl for 10 min and 3 washes with PBS. For the data in **Fig 1E-F, S1B, 3E-F** and **3E-F**, LDs were stained with MDH at RT for 10 min, followed by 3 washes with PBS. For the data in **Fig 3C-D, 4E-F** and nuclei were stained with DAPI for 10 min at RT and after 3 washes with PBS, LDs were stained with LD540 for 20 min at RT. LD dyes and DAPI were diluted in PBS. For the data in **Fig S3B-D**, samples were mounted with Mowiol-Dabco.

#### Fixed and live cell Airyscan imaging

Cells were imaged with Zeiss LSM 880 confocal microscope equipped with Airyscan (Fast) detector using a 63X Plan-Apochromat oil objective, NA 1.4. After fixing, cells were kept in PBS until imaging and stained for LDs immediately prior to imaging. Imaging was done in Airyscan super resolution mode. The Airyscan detector was adjusted regularly between acquisitions. For the data in **Fig 1E-H, S1B-D, 3A-B, 3E-F** and **3I-J**, z-stacks of cells were imaged using Airyscan microscopy. For the data in **Fig 2E**, single z-planes were captured in live cells. For the data in **Fig 1H**, fused cells containing both seipin-GFPx7 and seipin-SNAPf signal were imaged.

#### Fixed and live cell widefield imaging

Cells were imaged with Nikon Eclipse Ti-E microscope equipped with 60x Plan Apo λ oil objective, NA 1.4, and 1.5 zoom lens, Nikon Perfect Focus System 3, Hamamatsu Flash 4.0 V2 scientific CMOS camera and Okolab stage top incubator system. For the data in **Fig S2G**, a single z-plane was imaged every 30 min for 10 h. For the data in **Fig 3C-D, 4E-F** and **S3D**, z-stacks spanning the whole cell (step size 0.3 µm) were acquired.

#### Image processing

Airyscan images and videos were Airyscan-processed using the Zeiss Zen software package with identical (default) settings for all acquisitions. Deconvolution of widefield images was performed with Huygens software (Scientific Volume Imaging) using iterative Classic Maximum Likelihood Estimation. For downstream image analysis, z-stacks were maximum intensity projected using ImageJ FIJI or Matlab. Brightness, contrast and scale bars were adjusted in ImageJ FIJI and Corel Draw 2017 (64 bit).

#### Image analysis

For all LD analyses, except **Fig 3C-D**, LDs were segmented with Ilastik [51,52] using pixel and object classification mode, and final binary images were used for analysis. Subsequent cell segmentation was performed with CellProfiler [53], in a hierarchical manner, as previously reported [54]. For the data in **Fig 3C-D, 4E-F** and **S3I** cell nuclei were detected using the DAPI images with Otsu adaptive thresholding method. Touching nuclei were separated by built-in intensity methods. The cytoplasm was detected by utilizing the faint cellular background of DAPI channel, using intensity propagation based on Otsu adaptive thresholding method and the identified nuclei from the first step as a seed point. Finally, LDs were detected using static Otsu thresholding of the binary LD images generated in Ilastik or directly in CellProfiler (for the data in **Fig 3C-D** and **S3D)**, using CellProfiler module DetectAllDroplets [35]. Subsequent feature analysis, including LD size distributions and total LD numbers/cell was done with CellProfiler and custom Matlab software generated for post-processing reported in [35]. For the data in **Fig 1E-F**, images were manually cropped so that each image was from one cell, and separate cell segmentation was not necessary, and LD analysis was performed as described above. For the data in **Fig 3A-B, 3E-F** and **3I-J, S3F, S3J-K** each image was of a peripheral section of a single cell and cell boundary segmentation was performed in CellProfiler using faint background of antibody or LD staining. For the data in **Fig 3A-B, 3I-J, S3J-K** seipin and promethin foci were segmented with Ilastik using pixel and object classification mode and their overlap was analyzed using CellProfiler and custom Matlab software as described above. **Fig 3E-F** was analyzed similarly, except that promethin and LD overlap was analyzed. In all cases, structures were considered overlapping if they overlapped by at least 1 pixel. For the data in **Fig S3D**, cells were manually outlined and the mean intensity of promethin staining/cell (after background subtraction) was analyzed in ImageJ FIJI. For the data in **Fig 1H**, LDs were first segmented using Ilastik. Then, a static threshold was applied to the end-seipin-SNAPf channel and the sizes of thresholded LDs overlapping or not with seipin-SNAP foci were analyzed in ImageJ FIJI [55].

### Electron microscopy

Cells grown on fibronectin-coated coverslips were fixed with 2.5% glutaraldehyde in sodium cacodylate buffer, pH 7.4, supplemented with 2 mM CaCl2 for 25 min at RT. After washing, the cells were post-fixed with (non-reduced) 1% osmium tetroxide in 0.1 M sodium cacodylate buffer, pH 7.4, for 1 h at RT, dehydrated in ethanol series and flat embedded as described previously [56]. Ultrathin sections were post-stained with uranyl acetate and lead citrate, and imaged using a Jeol JEM-1400 microscope (Jeol Ltd) equipped with Orius SC 1000B camera (Gatan Inc.).

#### Electron tomography

For immuno-electron tomography (Fig 2H) the specimens were prepared as described in [34]. Briefly, after PLP-fixation [57] and permeabilization with 0.01% saponin, the cells were labeled with anti-GFP antibody for 1 h and nano-gold-conjugated anti-rabbit FAB-fragments for 1 h, post-fixed with 1% glutaraldehyde, and quenched with 50 mM glycine. Nano-gold particles were then intensified for 2 min using the HQ SILVER Enhancement kit followed by gold toning in subsequent incubations in 2% sodium acetate, 0.05% HAuCl4 and 0.3% sodium thiosulfate prior osmication, dehydration and flat embedding as described in [56]. A 230-nm-thick section cut parallel to the cover slip was subjected to electron tomography at nominal magnification of 19,000×, providing a 2×binned pixel size of 1.2 nm using Tecnai FEG 20 microscope (Thermo Fisher Scientific former FEI Corp.) operated at 200 kV. The alignment of the tilt series as well as reconstruction (SIRT technique using 14 iterations) were done with IMOD software package [58] (version 4.10.32). Segmentation and visualization were done using MIB [59] (version 2.601) and Amira (VSG, FEI Company) (version 5.3.2), respectively. The immunolabeling is not shown, as the labeling was clearly denser at the outskirts and edges of the S166D-induced ER sacs, suggesting that steric hindrance may prevent diffusion of the antibodies to these structures.

### Computational methods

#### Protein modeling

Seipin oligomer was modeled as an all-atom description using the cryo-EM structure of human seipin (PDB ID: 6DS5 [31]). The structure of an individual seipin monomer comprised of amino acids (aa.) 25-263 corresponds to the N-terminal transmembrane helix (27-47), topological domain (48-242), and the C-terminal transmembrane helix (243-263). The terminal topological domains (aa. 1-24 and 264-398) were not modeled into the seipin structure. The TMDs of the protein (corresponding to aa. 25-47 and 243-263) and the missing segments of the lumenal domain (corresponding to aa. 48-59 and 220-242) were built into the cryo-EM structure using the PyMol software. The transmembrane region of the protein was modeled as a regular right-handed alpha-helix. Residue N88 was N-glycosylated (Manα1-6(Manα1-3)Manβ1-4GlcNAcβ1-4GlcNAcβ1-Asn).

#### Building the simulation system

Continuing at atomic resolution, a multi-component lipid bilayer (**Table 1**) mimicking the lipid profile found in the ER [60] was constructed around the seipin oligomer. All lipids were randomly distributed in the membrane plane. The transmembrane distribution of TAGs was symmetric. For other lipids, the transmembrane lipid distribution in the ER is not well known. Some studies have reported that it is asymmetric [61,62], while there are also views in favor of a symmetric distribution [63]. In the present work, for the sake of completeness, we considered both options: an asymmetric transmembrane lipid distribution that was tuned from previous simulation studies [64] and a symmetric distribution. The results presented in this article are based on the asymmetric distribution, unless otherwise noted, but the results based on a symmetric distribution are consistent with those of the asymmetric distribution. Lateral pressure was applied to ensure proper packing of the lipids around the oligomer structure. The TMDs and the α2-α3 helical segment (aa. 154-170) of the protein were embedded within the bilayer. The system was solvated (full hydration) and counter ions were added to neutralize the system. Additionally, 0.15 M of KCL was used to mimic the physiological salt concentration in the cytosol.

For TAGs, the objective was to track non-equilibrium diffusion and clustering processes where initially random distributions of TAGs self-assemble spontaneously to form TAG-rich clusters. Given this, the simulation systems equilibrated during the clustering processes.

#### Simulation parameters

Simulations of both all-atom and coarse-grained model systems (**Table 2**) were carried out using the GROMACS simulation package, version 2016.4 [65]. In all-atom simulations, we used the CHARMM36 force field [66–69] for the protein, lipids, ions, and TAGs. For water, the force field used was CHARMM TIP3P. The simulation systems were first energy minimized and pre-equilibrated under constant temperature and pressure for 1 ns. Temperature was maintained at 310 K using the Nose-Hoover thermostat [70] with a coupling constant of 0.5 ps. Pressure was maintained semi-isotropically at 1 atm using the Parrinello-Rahman barostat [71] with a coupling constant of 2 ps. The time-step used for integrating the equations of motion was 2 fs. The neighbor list was updated every 10 steps using the Verlet cut-off scheme. The LINCS algorithm [72] was used to constrain the motion of covalently bonded hydrogen bonds. Short-range interactions (electrostatic and van der Waals) were cut-off at 1.0 nm. The long-range component of electrostatic interactions was calculated using the Particle Mesh Ewald (PME) method [73]. For every system, production simulations of 1 μs length were performed in two replicates.

**Table 2:** Systems explored through atomistic and coarse-grained simulation models.

| Simulation | Seipin | Membrane system<br>(mol% TAG) | Simulation<br>time (in μs) | No. of<br>simulations |
| --- | --- | --- | --- | --- |
| Atomistic | Wild-type | 2.5 (randomly distributed) | ~1 | 2 |
|  | Wild-type | 2.5 (clustered within lumen) | ~1 | 2 |
|  | Mutant (S166D) | 2.5 (randomly distributed) | ~1 | 2 |
|  | Mutant (S166D) | 2.5 (clustered within lumen) | ~1 | 2 |
|  | Mutant (S166D) protonated | 2.5 (randomly distributed) | 0.2 | 2 |
|  | Mutant (S166A) | 2.5 (clustered within lumen) | ~1 | 2 |
|  | Mutant (S165A-S166A) | 2.5 (clustered within lumen) | ~1 | 2 |
| Coarse-grained | No seipin | 2.5 | 5 or 30 | 10 |
|  | Wild-type | 2.5 | 5 or 30 | 10 |
|  | No seipin | 4.85 | 5 | 10 |
|  | Wild-type | 4.85 | 5 | 10 |
|  | Mutant (S166D) | 4.85 | 5 | 10 |
|  | Mutant (S166A) | 4.85 | 5 | 10 |
|  | Mutant (S165A-S166A) | 4.85 | 5 | 10 |
|  | No seipin | 1.25 | 5 | 10 |
|  | Wild-type | 1.25 | 5 | 10 |
|  | Mutant (S166D) | 1.25 | 5 | 10 |
|  | Mutant (S166A) | 1.25 | 5 | 10 |
|  | Mutant (S165A-S166A) | 1.25 | 5 | 10 |
| | β2-adrenergic receptor | 1.25 | 5 | 10 |
|  | Sigma-1 receptor | 1.25 | 5 | 10 |
|  | Seipin luminal domain only | 1.25 | 5 | 10 |

In coarse-grained simulations, a model consistent with the MARTINI framework (version 2.2) [72,74,75] was first generated from the atomistic protein model using the martinize.py script (http://cgmartini.nl/index.php/tools2/proteins-and-bilayers). The residue N88 was not glycosylated in the coarse-grained model of seipin. The systems were again first energy minimized and pre-equilibrated under constant temperature and pressure for 10 ns. Temperature was maintained at 310 K by the V-rescale thermostat [76] and pressure was maintained semi-isotropically at 1 atm using the Parrinello-Rahman barostat. The standard MARTINI water model was used with a relative dielectric constant of 15. A cut-off of 1.1 nm was used for Coulombic and van der Waals interactions. The integration time step was chosen as 20 fs. Production simulations of 5 μs each were carried out in 10 replicates starting from different initial conditions. Further, to foster the sampling of clustering of TAGs at longer time scales, three sets from each of the coarse-grained systems with a TAG concentration of 2.5 mol% were randomly chosen and extended to 30 μs.

The simulations of seipin were complemented by coarse-grained simulations of other membrane proteins that do not play a known role in LD biogenesis: ß2-adrenergic receptor (B2AR) (PDB ID: 3D4S [77]) and sigma-1 receptor (S1R) (PDB ID: 5HK1 [78]). The simulation protocol was the same as that used for seipin.

#### Analysis of simulation data

All analysis was carried out using standard GROMACS tools. The mean curvature of the leaflets was calculated using g_lomepro [79]. VMD [80] was used for visualizing the simulations and rendering images.

Average TAG occupancies were calculated for each amino acid residue in the seipin oligomer **(Fig 1C)** as follows: For each ns, the smallest distance between the selected amino acid residue and all TAG molecules in the membrane was computed. The residue was considered to interact with TAG if the smallest distance was 0.5 nm or less. This analysis was carried out separately for 0-1 and 4-5 µs of simulation time, over all simulation repeats of the same system, and separately for every residue of seipin, thus giving an average occupancy number for each residue. Occupancy value one indicates that the selected amino acid residue always interacted with TAG, while a value of zero indicates that the amino acid residue never experienced interactions with TAG.

To calculate the fraction of TAGs clustered in a bilayer **(Fig 4B, D)**, we calculated the smallest distance between a chosen TAG and the rest of the TAG molecules. The chosen TAG molecule was considered to interact with other TAGs if the smallest distance was 0.5 nm or less, and was considered to be part of a cluster if it interacted with at least 2 other TAG molecules. Therefore, the fraction of TAGs clustered in a bilayer exclude monomeric and dimeric TAGs.

Short-time diffusion coefficient *D* for TAGs inside the seipin complex **(Fig 2A)** was calculated using a technique discussed by Vuorela et al. [81]. For a given molecule, one records the 2D (in the plane of the membrane) displacement (Δ*r*) during a time interval Δ*t*. When the molecule diffuses over a long period of time, one finds the distribution of these displacements, *P*(Δ*r*), which can be fitted to the theoretical distribution [81]. The fitting yields a value for the short-time diffusion coefficient *D*. Since the analysis was made over a period of Δ*t* = 1 ns, this analysis yields a diffusion coefficient that characterizes diffusive behavior in the nanometer-scale, and is thus able to document TAG binding for S166 and its mutants.

#### Statistics

Statistical analysis was performed with Graphpad Prism (Graphpad Software), using statistical tests as indicated in the figure legends. p-values < 0.05 were considered significant. Throughout the figure legends: * p <0.05, ** p <0.005, *** p < 0.0005.

## Acknowledgements

We thank Anna Uro and Suvi Saarnio for excellent technical assistance. We thank Helsinki Institute for Life Science (HiLIFE) Light and Electron microscopy platforms and Flow cytometry unit and Biocenter Finland for infrastructure support. This study was financially supported by the Academy of Finland Center of Excellence Program (grant 307415 to EI and IV), Academy of Finland grants (282192 and 312491 to EI; 287975 to EJ), Sigrid Juselius Foundation (EI, IV), Jane and Aatos Erkko Foundation (EI) and the HiLIFE Fellow Program (EI, IV, EJ); VS acknowledges support from the Doctoral Progamme in Biomedicine, Finnish Medical, Paulo, Alfred Kordelin, Maud Kuistila, Biomedicum Helsinki and Emil Aaltonen Foundations. We thank CSC – IT Center for Science Ltd. (Espoo, Finland) and the Finnish Grid and Cloud Infrastructure (FGCI) for computing resources.

## Author contributions

EI, IV, XP and VTS designed the study. XP performed and analyzed MD simulations, with help from IV. VTS performed and analyzed cell experiments, with help from SL and KV. HV and EJ performed EM. VTS and EI wrote the manuscript and all authors commented on the manuscript.

## Supplementary figure legends

**Fig S1.**
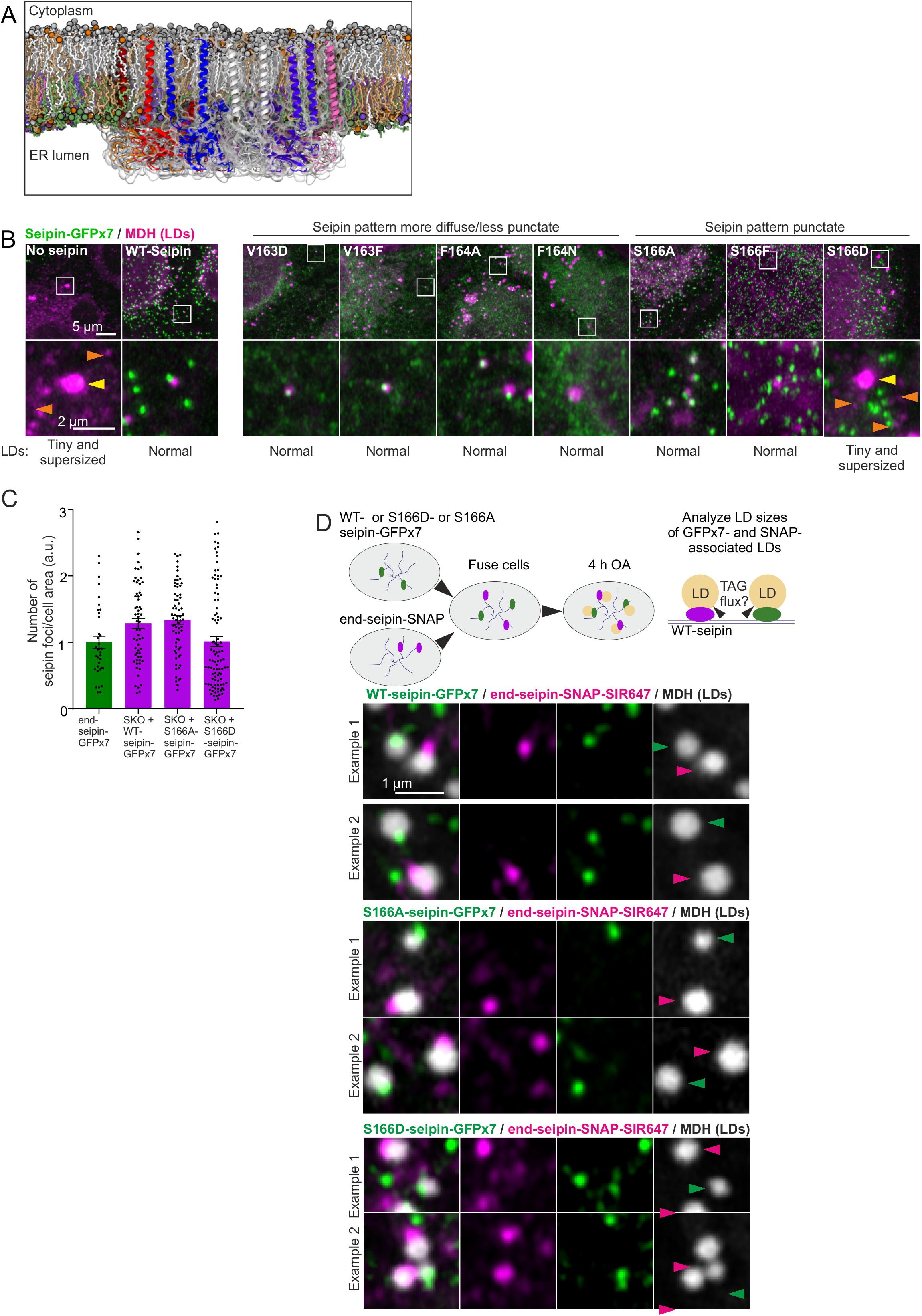
**A)** Side view of the seipin oligomer in the model ER bilayer. The model was based on the cryo-EM structure of human seipin and the TMDs were modeled. The lipid composition was modeled to match that reported for the ER. **B)** A431 SKO cells stably expressing the indicated plasmids were delipidated for 3 days and treated with 200 µM OA for 1 h. Cells were fixed, LDs stained, and imaged by Airyscan microscopy. Orange arrowheads: tiny LDs, yellow arrowheads: supersized LDs. **C)** Related to Fig 1E-F. Indicated cell lines were delipidated for 3 days and treated with OA for 1 h. Cells were fixed, LDs stained, and imaged by Airyscan microscopy. The number of seipin foci/cell area was analyzed and plotted. n = 35 cells for end-seipin-GFPx7, >60 cells/group for the others, 3-4 experiments. **D)** Exemplary crops of nearby LDs from cells treated as indicated. Green arrowheads: WT-, S166A- or S166D-seipin-GFP associated LDs, magenta arrowheads: end-seipin-SNAP associated LDs. See Fig 1H for analysis of the data.

**Fig S2.**
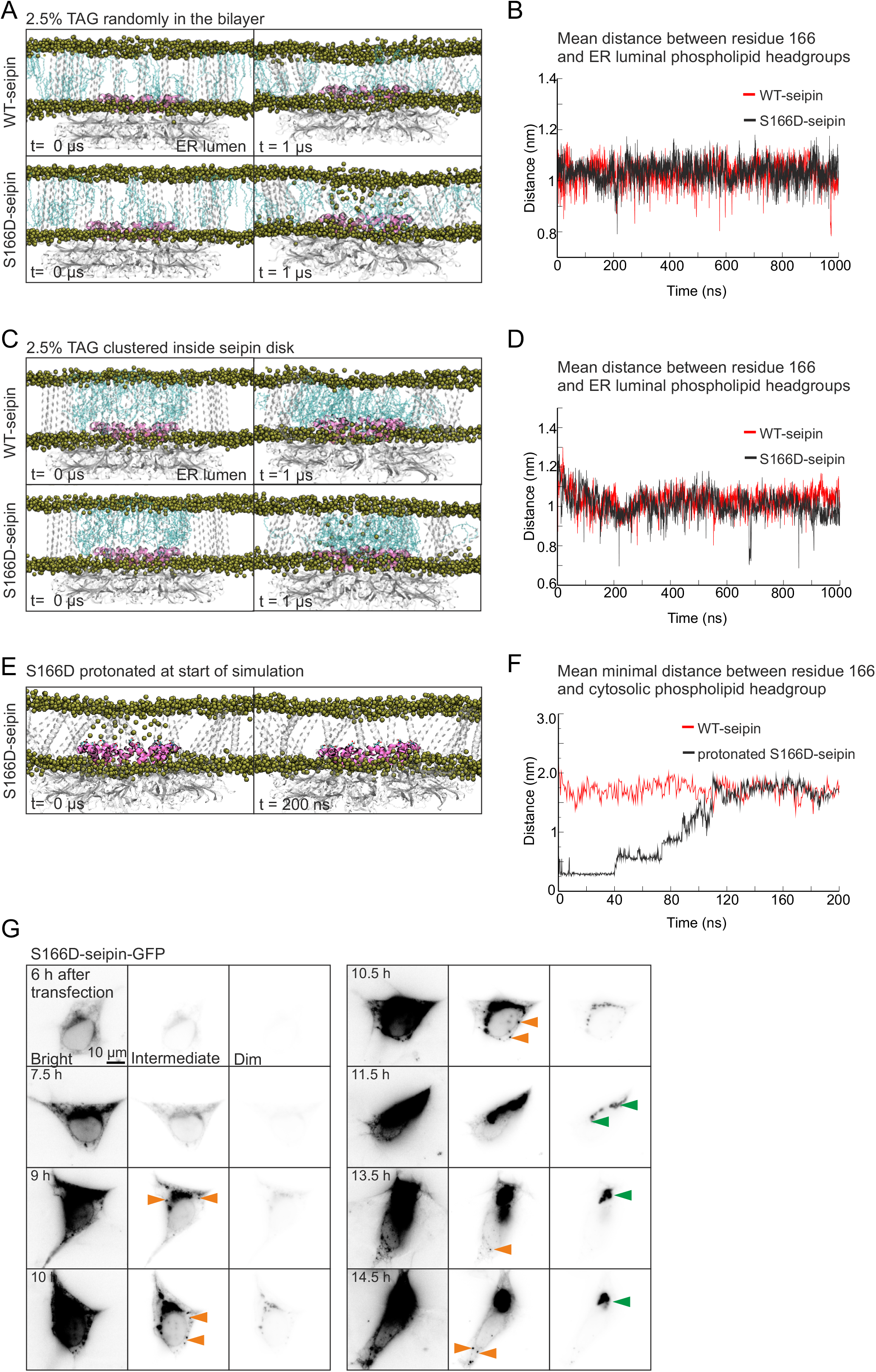
**A)** Snapshots of atomistic simulations showing the relative position of α2-α3 helices (pink) with respect to the head group region of the PLs in the bilayer (represented by phosphate atoms shown as yellow spheres) at the beginning (0 µs) and end (1 µs) of simulation. The system consists of 2.5 mol% TAG randomly distributed in the bilayer around the seipin oligomer. The TM helices and luminal domain are shown in white (transparent). The TAGs are shown in cyan (transparent). **B)** Center of mass distance between the residue 166 (all) and the phosphate atom of PLs in the luminal leaflet of the bilayer over the simulation period for the systems shown in A. **C)** Snapshots of atomistic simulations showing the relative position of α2-α3 helix with respect to the head group region of the PLs in the bilayer at the beginning (0 µs) and end (1 µs) of simulation. The system consists of 2.5 mol% TAG clustered within the lumen of the seipin oligomer. Coloring as in A. **D)** Center of mass distance between residue 166 (all) and phosphate atom of PLs in the luminal leaflet of the bilayer over the simulation period for the systems shown in C. **E)** Snapshots of atomistic simulations. Left snapshot shows the membrane deformation (depicted by the invagination of phosphate atoms from the cytosolic leaflet into the bilayer) in the S166D-seipin system. All S166D residues were then protonated (0 µs) and simulated. Right snapshot at the end of the 200 ns simulation shows that the deformation has relaxed. Coloring as in A. **F)** Minimum distance between the protonated S166D residue 166 (all) and phosphate atoms of the PLs in the cytosolic leaflet of the bilayer over a simulation period of 200 ns. Similar data for wild-type S166 residue is shown for reference. **G)** HEK 293A cells were transfected with S166D-seipin-GFP and imaged live by widefield microscopy. A single cell is shown over time, orange arrowheads indicate smaller GFP foci, which upon rising fluorescence levels coalesce to a larger membranous aggregate (green arrowheads).

**Fig S3.**
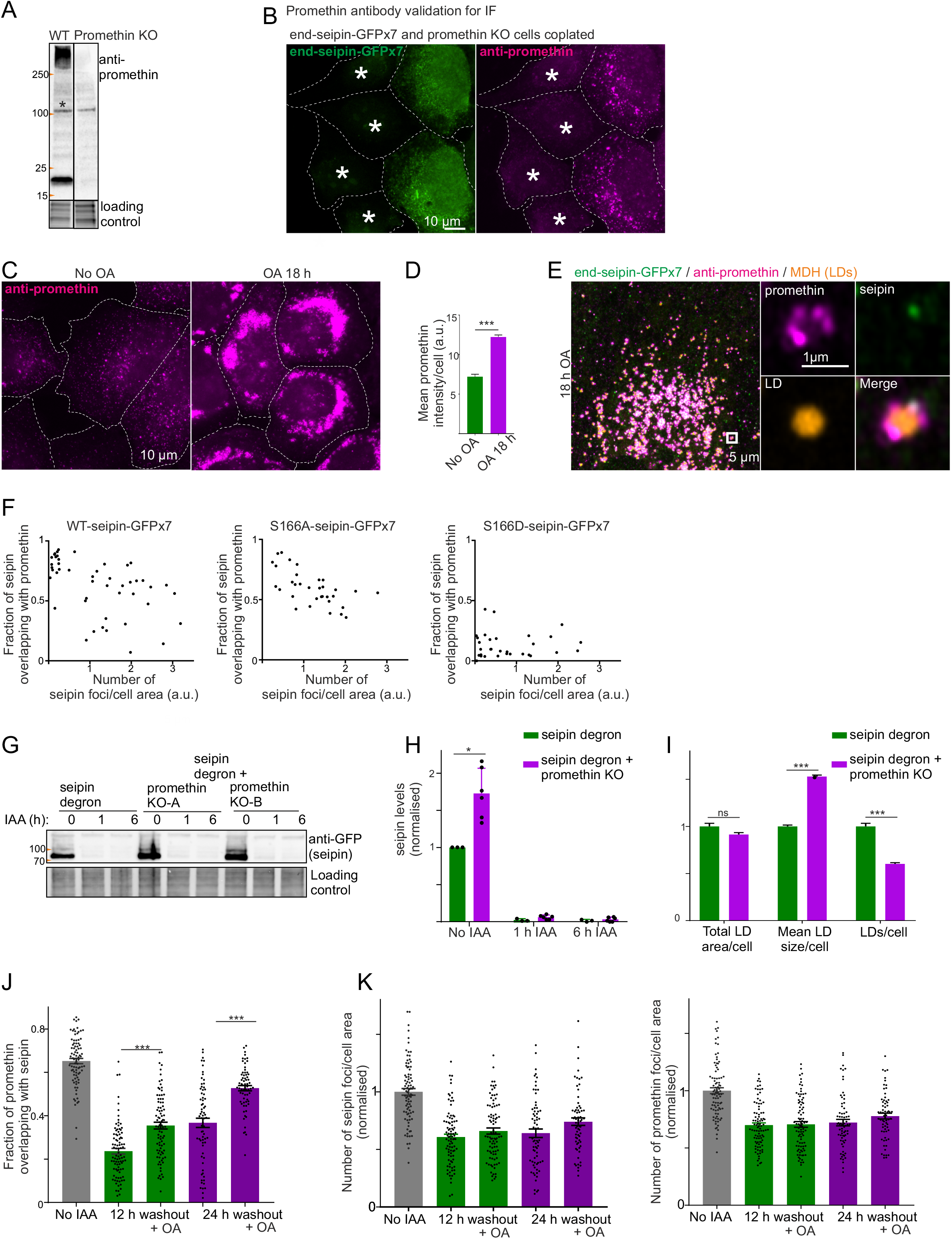
**A)** Representative immunoblot of WT A431 cells and promethin KO cell pool using anti-promethin antibody. Asterisk indicates unspecific band. Note both monomer-sized (17 kDa) and larger (>250 kDa) specific bands. **B)** end-seipin-GFPx7 cells and promethin KO cells were coplated, fixed and stained with anti-promethin antibodies. Asterisks indicate promethin KO cells that have not been engineered to harbor seipin-GFPx7. Note bright promethin staining in the end-seipin-GFPx7 cells. Maximum projections of widefield z-stacks. **C)** Cells were cultured in complete medium or additionally treated with 200 µM OA for 18 h, fixed and stained with anti-promethin antibodies. **D)** Analysis of C. Bars = mean +/- SEM, n = 71-180 cells/group, representative experiment repeated once with similar results. Statistics: Mann-Whitney test. **E)** end-seipin-GFPx7 cells were treated with 200 µM OA for 18 h, fixed and stained with anti-promethin antibodies and MDH. Maximum projection of two Airyscan z-slices 210 nm apart. **F)** In relation to Fig 3B, the fraction of seipin foci overlapping with promethin foci are plotted relative to the number of seipin foci/cell area. A value of 1 in number of seipin foci/cell area corresponds to the mean seipin foci/cell area detected in end-seipin-GFPx7 cells in Fig S1C. **G)** Immunoblots of seipin degron cells with or without promethin KO treated with IAA. **H)** Analysis of F). Bars = mean +/- SEM, n = 3-6 replicates/group, 2 experiments. Promethin KO data are pooled from KO-A and KO-B pools. Statistics: Mann-Whitney test. **I)** In relation to Fig 3C-D, an additional representation of the “No IAA” data of that panel. Seipin degron cells +/- promethin KO were delipidated for 3 days and treated with OA for 1 h as in Fig 3C and LDs analyzed. Bars = mean +/- SEM, n > 500 cells/group, 3 experiments. Promethin KO data are pooled from KO-A and KO-B pools. Statistics: Mann-Whitney test. **J)** Additional analysis of the data in Fig 3I-J. Cells were treated as in Fig 3I and the fraction of promethin foci overlapping with seipin foci was analyzed. Bars = mean +/- SEM, n = >60 cells/group, 4 experiments. Statistics: Kruskal-Wallis test followed by Dunn’s test. **K)** Additional analysis of the data in Fig 3I-J. The number of seipin or promethin foci/cell area is plotted, normalized to the number of seipins or promethins/cell area in the “No IAA” sample. Bars = mean +/- SEM, n = >60 cells/group, 4 experiments.

**Fig S4.**
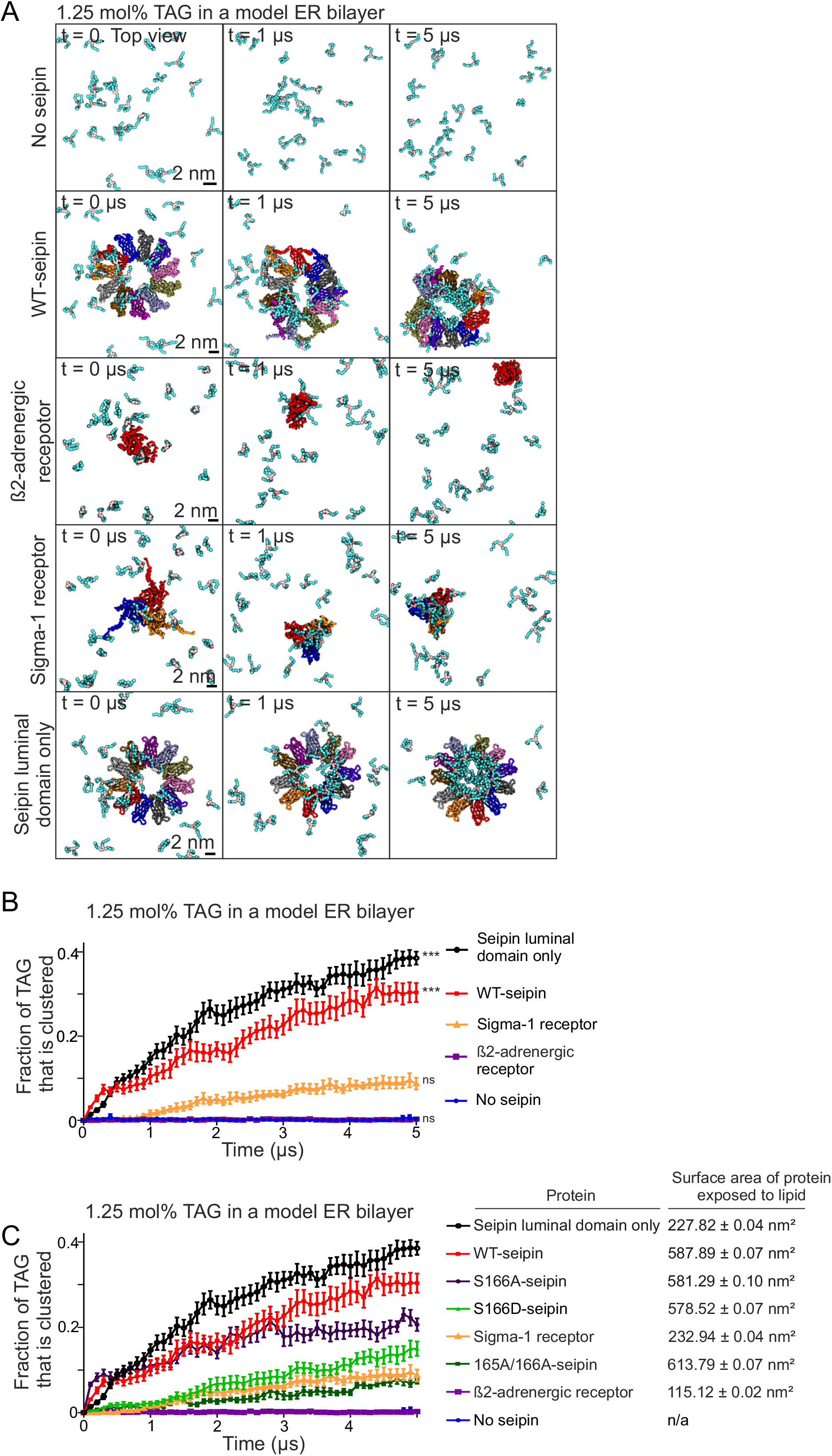
**A)** In relation to Fig 4A. Snapshots of coarse-grained simulations with 1.25 mol% TAG in the bilayer. Results are shown for two additional integral membrane proteins to demonstrate their TAG affinities. The color coding is as in Fig 1A. Snapshots of “No seipin” and “WT seipin” are the same as in Fig 4A. **B)** Analysis of A. Data points: mean, +/- SEM, n = 10 simulations/system. Data for “No seipin” and “WT seipin” are same as in Fig 4B. Statistics are based on the final time points of analysis using Kruskal-Wallis test followed by Dunn’s test, comparing against no seipin. **C)** Additional analysis of the data in Fig 4B and S4B. Total surface area of protein exposed to membrane lipids was calculated using GROMACS tool sasa. Data points: mean, +/- SEM, n = 10 simulations/system. Whilst membrane proteins are expected to reduce the mobility of nearby lipids relative to the membrane exposed protein surface area (s.c. blocking effect), this does not explain the TAG clustering effect of WT seipin.

